# Qualitative and quantitative analysis of the constitutive bark of *Q. ilex* x *Q. suber* hybrids

**DOI:** 10.1101/2022.07.12.499723

**Authors:** Gonzalo de Burgos, Eduardo Díez-Morales, Unai López de Heredia, Álvaro Soto

## Abstract

Hybridization and introgression between cork oak (*Quercus suber*) and holm oak (*Q. ilex*) have traditionally been reckoned as undesirable processes, since hybrid individuals lack the profitable bark characteristics of cork oak. Nevertheless, a systematic and quantitative description of the bark of these hybrids at the microscopic level based on a significant number of individuals was not available to date.

In this work we provide such a qualitative and quantitative description, identifying the most relevant variables for their classification. Hybrids show certain features intermediate between those of the parent species, as well as other unique features, as the general suberization of inactive phloem, reported here for the first time. These results suggest a relevant hybridization-induced modification of the genetic expression patterns. Therefore, hybrid individuals provide a valuable material to disentangle the molecular mechanisms underpinning bark development in angiosperms.

## Introduction

Cork oak (*Quercus suber* L.) and holm oak (*Q. ilex* L.) are two ecologically, economically, environmentally and socially relevant oak species that are key elements in open woodlands from the western Mediterranean basin. Both species differ in morphological traits of bark, cupules of the acorns, or leaves (Amaral Franco 1990) and they belong to two different subsections (Ilex and Cerris) within section Cerris (McVay et al. 2017; Denk et al. 2017).

Possibly, bark anatomy is the most discriminating phenotypic character between cork and holm oaks at the adult stage. The bark of most Euromediterranean oak species is made up of a rhytidome, a structure formed by successive phellogens that produce thin, suberized and intricate phellem layers enclosing heterogeneous cortical tissues (parenchyma, fibers, etc.) and collapsed phloem cells. This type of outer bark has been described in detail for *Q. robur* (Trockenbrodt 1991), *Q. petraea* (Gricar et al. 2015), *Q. faginea* (Quilhó et al. 2013), *Q. cerris* (Sen et al. 2011) and *Q. ilex* ssp. *rotundifolia* (Sousa et al. 2021).

In contrast, the outer bark of *Q. suber* is thought to be caused by the differentiation of a single long-living phellogen that generates new layers of suberized and sometimes lignified cells each year, rather than producing a rhytidome (Graça and Pereira 2004). This bark differentiation pattern results in cork, a thick regular suberized material with extraordinary protective and insulation properties against pathogen attacks or fire (Pereira 2007; Pausas et al. 2009). Indeed, cork is exploited in the Mediterranean countries due to its renewable character and high economic and social value in rural areas. Cork is mainly formed by dead, empty prismatic cells that are stacked by their bases in radially aligned rows disposed in parallel without intercellular spaces (Leite and Pereira 2017). The cork cell walls are covered with suberin, an inert hydrophobic substance partially similar to lignin and cutin. Virgin cork is more irregular and presents more lignified cells than the traumatic cork that is produced after successive peeling, which is the basis of commercial cork exploitation (Pereira 2007).

Although *Q. suber* and *Q. ilex* have slightly different ecological constraints, they frequently co-exist forming mixed stands (particularly with ssp. *rotundifolia*) where they are able to outcross to produce fertile hybrids (López de Heredia et al. 2017). The occurrence of *Q. ilex x suber* hybrids (*Quercus x morisii* Borzí; *Quercus x avellaniformis* Colmeiro & E. Boutelou) has long been known and botanical descriptions present these hybrids as having viable acorns with conical cupules with free bracts, glabrescent leaves of light green tone and strongly cracked bark (Colmeiro and Boutelou 1854; Laguna 1881; Borzi 1881; Natividade 1936; Camus 1936– 1954). These hybrids also present micro-morphological and anatomical characters related to the presence/absence and distribution of foliar trichomes (Vázquez 2015) and to the thickness of the leaf lamina (López de Heredia et al. 2018).

Traditionally, it has been considered that hybridization produces bad quality cork and that first generation hybrids are not suitable for cork extraction (Varela et al. 2008). Early qualitative studies of the bark anatomy of *Q. ilex x suber* hybrids reported high variability and described some hybrids showing corky barks, others resembling the rhytidome structure of *Q. ilex*, but with thicker layers of phellem, and some others presenting intermediate traits (Natividade 1936). However, a robust quantitative description of the anatomical features of the bark of hybrids is lacking. To our knowledge, this issue has only been afforded from a qualitative perspective using a limited number of putative hybrids (Natividade et al. 1936; Llensa de Gelcén 1943) whose precise level of hybridization could not be checked with molecular markers (Soto et al. 2018).

In the present paper, we adopt a qualitative and quantitative approach to assess bark anatomical features of a higher number of adult *Q. ilex x suber* hybrids of known hybridization level inferred with high resolution molecular markers (López de Heredia et al. 2020).

## Material and Methods

### Sampling and laboratory procedures

Twenty naturally grown hybrid trees were identified and sampled in a mixed holm oak-cork oak forest in Fregenal de la Sierra (Extremadura, SW Spain). Hybrids were initially identified according to morphological characters described by Natividade (1936) and further confirmed with molecular markers (López de Heredia et al. 2020). Samples were also collected from one cork oak (*Q. suber*) and two holm oak (*Q. ilex* ssp. *rotundifolia*) trees as a reference, since previous works have reported almost no variability among individuals within these species (Sousa et al. 2021). Outer bark samples (5×12 cm) were harvested from the south-facing part of the trees at DBH (1.30 m) using a hammer and a chisel.

Six cubical prisms (2 × 4 cm) were extracted from each sampled bark piece. After polishing the cube surfaces, samples were submerged in 70% ethanol and surface photographs were taken with a Moticam 1080 camera on a Nikon 8MZ-2T binocular lens. In addition, 25 μm thick cross-sections were obtained with a Leica SM2400 microtome. Cross-sections were heated in a sodium hypochlorite solution prior to staining for suberized and lignified tissue observation using an Olympus BX51 microscope equipped with a fluorescence lamp Olympus U-RFL-T (excitation at 340-380 nm and 410-450 nm barrier filters) and a Moticam 1080 camera (Motic Microscopy). The phloroglucinol-HCl test (Nêmec 1962) was performed for visualization of lignified and suberized cells under tungsten and UV light. A drop of 1% phloroglucinol:ethanol solution (w/v) was poured on washed crossed-sections, followed by the addition of 25 μl of 35% HCl. Using this procedure lignin appears stained in red, while quenching of lignin autofluorescence under UV light by phloroglucinol-HCl allows the identification of suberized tissue (Briggs 1985).

### Quantitative analysis

The obtained photographs were treated with the Motic Images Plus 2.0 (Motic Microscopy) and ImageJ (https://imagej.nih.gov/ij/index.html) software applications to measure seven bark quantitative and two qualitative categorical traits (**Table 1**) based on a preliminary exploratory analysis of the most relevant traits for outer bark characterization in cork oak, holm oak and their hybrids (López de Heredia et al. 2017). In total, 407 pairs of photographs obtained with visible light and ultraviolet light were measured.

**Table 1:** Description of the measured traits for outer bark anatomical characterization. VL: visible light; UV: Ultra-violet light

| **Trait** | **Description** | **Microscope**<br>**light exposure** |
| --- | --- | --- |
| PPH | Percentage of ordered suberized cells<br>(phellem cells) | UV |
| PT | Mean phellem cell number (4 random<br>measurements) | UV |
| Nmax | Maximum number of phellem cells | UV |
| Nmin | Minimum number of phellem cells | UV |
| PHT | 1: Rhytidome with non-functional phloem<br>2: Rhytidome with massive phellem | Both |
| PP | Percentage of parenchymatic cells | Both |
| PS | Percentage of schlerenchymatic cells | VL |
| SG | 0: lack of schlerenchymatic cells<br>1: isolated schlerenchymatic cells<br>2: grouped schlerenchymatic cells | VL |
| DS | Percentage of disordered suberized tissue | UV |

A categorical variable (PHT) evaluated *de visu* the type of bark in the sample (1: rhytidome with non-functional phloem; 2: rhytidome with massive phellem cells). Phellem related quantitative traits scored the percentage of phellem cells in the sample (PPH), the mean thickness of phellem tissue from measures in four random points of the photograph (PT) and the maximum and minimum number of phellem cells (Nmax and Nmin) in each sample. Cork oak presents a massive phellem that occupies the entire photograph; therefore, an arbitrary PT=100 cells was established to allow the comparison with the phellem thickness of the rhytidomes of holm oak and the hybrids. The percentage of non-functional phloem tissue between successive periderms (PP) was estimated using both visible and UV lights. Photographs taken under visible light were also employed to ascertain the percentage of lignified schlerenchymatic cells (PS) and their spatial arrangement through a categorical variable (SG), as they may occur isolated (1), forming groups of lignified cells (2), or not occur at all (0). Finally, providing that some of the hybrid samples showed disordered suberized tissue, we calculated the percentage of this type of cells in the sample (DS).

Median, mean, maximum/minimum values and standard deviation were scored for each quantitative variable at individual tree and species (cork oak, holm oak and hybrids) levels, respectively. To identify the most significant variables discriminating between bark types, ANOVAs were performed for each quantitative variable, according to the following model:

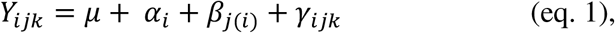

where μ is the mean value of the variable Y, α_i_ is the effect of species *i*, β_j(i)_ is the effect of the individual *j* within species *i* and γ_ijk_ is the error term for the observation *k* of the individual *j* within species *i*. The differences between species and individuals for the categorical variables PHT and SG were evaluated using a Kruskal-Wallis test.

A principal component analysis (PCA) was performed by scaling the original variables with the “prcomp” function from the R 4.0.3 “stats” package (R Core Team 2020). The two principal components (PCs) explaining most of the total variance were selected. New variables were created from the scores of the components, thus simplifying further analysis. The mean scores by individual of the first two principal components were bi-plotted to graphically inspect differences between species, and the correlation of these values with the introgression level of the hybrid trees estimated by López de Heredia et al. (2020). The introgression level was estimated using the q_s_ coefficient provided by a Bayesian approach implemented in STRUCTURE v. 2.3.4 (Pritchard et al., 2000) for each individual that may show values between 0 and 1. q_s_ values close to 0 suggest ilex-like trees, q_s_ values close to 1 suggest suber-like individuals and q_s_ values close to 0.5 suggest the tree is a first generation (F1) hybrid.

## Results and discussion

Our observations of *Q. suber* samples are consistent with the descriptions reported in previous works (f.i., Natividade 1936; Pereira et al. 1987). The outer bark of this species is formed by a single periderm, several centimeters thick, and with deep cracks and furrows, mainly longitudinal, not reaching the phellogen. This periderm appears, in a transverse section, as a light colored layer, where annual growth rings can be distinguished as darker lines. Microscopically the periderm appears formed mainly by empty, suberized cells, with few schlerenchymatic nodules (**Figure 1**).

**Figure 1:**
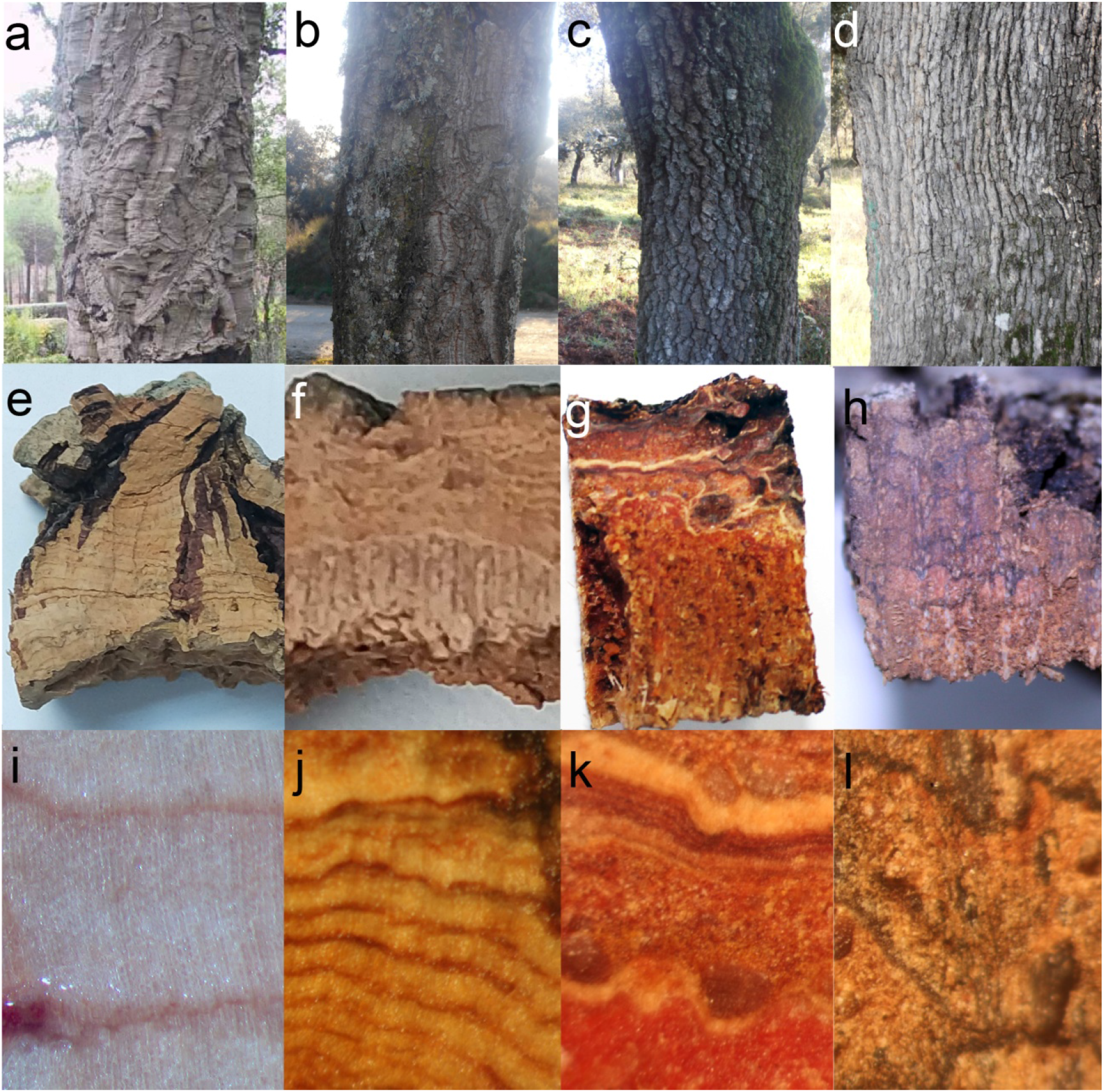
Barks of *Q. suber* (a, e, i), suber-like hybrid (b, f, j), ilex-like hybrid (c, g, k) and *Q. ilex* ssp. *rotundifolia* (d, h, l) as seen in the tree (a-d), in transverse section, both *de visu* (e-h) and under magnifying glass (i-l).

On the contrary, *Q. ilex*, as most species of the genus, shows a rhytidome, with several successive anastomosed periderms and lignified inactive phloem in between. Macroscopically, holm oak outer bark is finely crackled in small tiles, which eventually fall from the tree. In a transverse section, periderms can be distinguished as dark, thin lines; inactive phloem presents a brown-reddish color, with whitish multiseriate lignified rays and schlerenchymatic nodules, consistently with the observations by Sousa et al. (2021). As seen under the microscope, periderms are few cells thick, and become more abundant in the outer part of the bark, as mechanical tensions induce the differentiation of new phellogens in the inactive phloem comprised between periderms.

Much larger variability can be observed in hybrid individuals. Externally, most of them show an intermediate aspect, with deeper cracks than *Q. ilex* bark, and larger tiles. Some individuals present longitudinal furrows more similar to those of *Q. suber*, and with orange color in the inner part. In transverse sections most individuals show abundant light-colored layers, corresponding to periderms. Very often they appear arranged in parallel, although in other cases they look anastomosed. Abundant whitish multiseriate rays and schlerenchymatic nodules are also visible within inactive phloem. Under the microscope, most individuals show very abundant periderms, often much thicker than in *Q. ilex*. These periderms also appear much closer one to another than in holm oak. More strikingly, most individuals also present suberization in inactive phloem cells (**Figure 2**). This feature has not been observed in *Q. suber* or *Q. ilex* samples or in any other species, to our knowledge.

**Figure 2:**
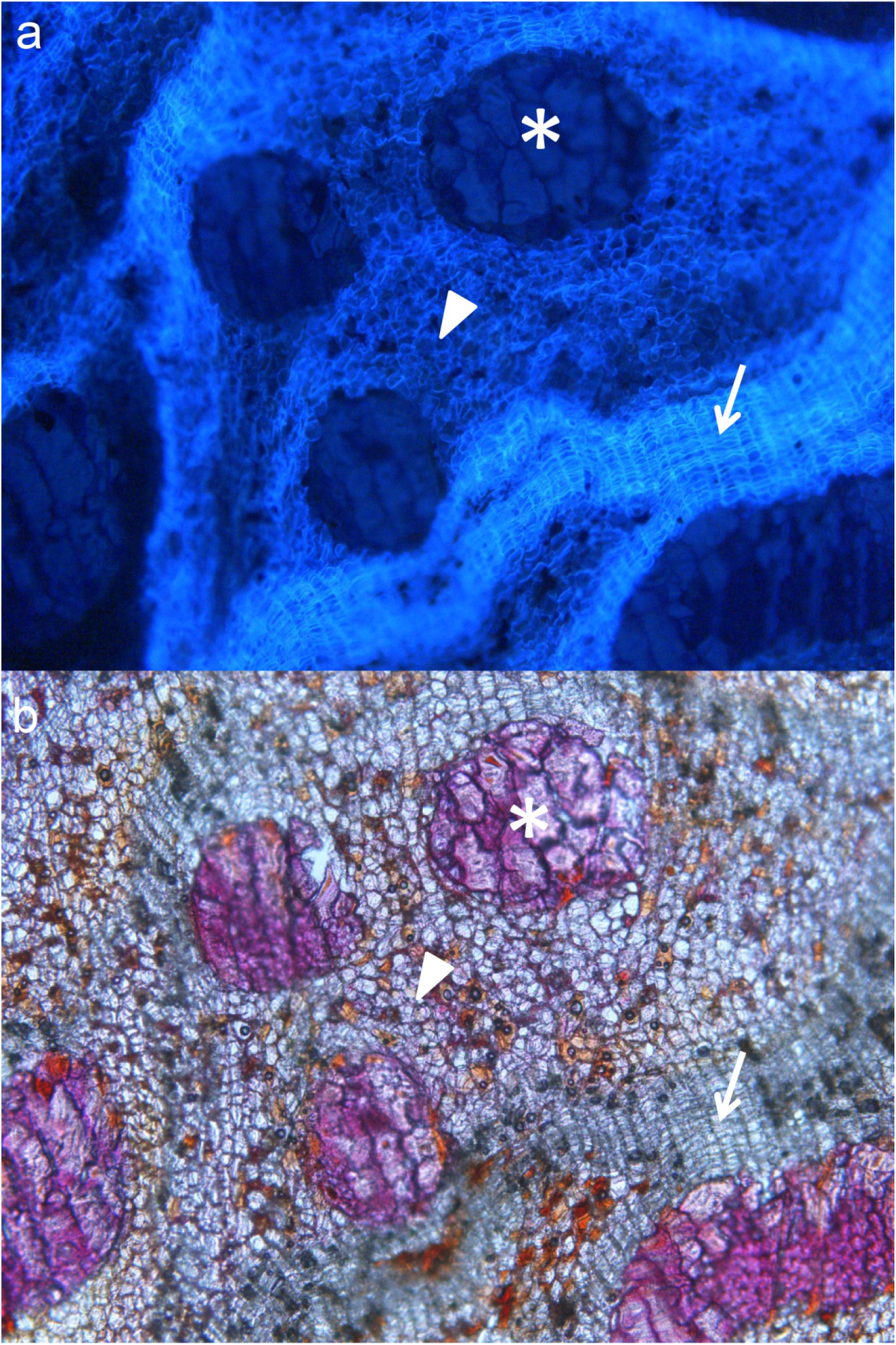
Cross section of the bark of a hybrid tree under UV (a) and visible light (b). Periderms (arrow), schlerenchymatic nodules (asterisk), and a high proportion of suberized inactive phloem (arrowhead) can be seen.

Once performed these explorative observations, we defined a series of qualitative and, mostly, quantitative parameters or variables in order to perform a statistical comparison of the barks (**Table 1**). Measurements of these variables show a great variability between species as well as among hybrid trees (**Figure 3**; **Table S1**). Actually, within-individual variability was also detected in hybrid trees, with some individuals showing areas with thicker phellems and other areas with higher presence of inactive phloem and thinner periderms. Values observed for parental species are consistent with those reported previously. Thus, *Q. suber* samples are characterized by a very thick periderm, accounting for most of the outer bark, with a large, indefinite number of phellem cells (set to 100 in order to ease further numerical analysis), with a low proportion of schlerenchymatic cells. On their side, a larger proportion of non-suberized inactive phloem (almost 70% of their outer bark) characterizes *Q. ilex* samples, with a significant proportion of lignified rays and schlerenchymatic nodules (9-14%). Phellem tissue accounts for approximately 20% of the bark transverse area, and is comprised of several thin (∼10 cells thick), anastomosed periderms, which is consistent with previous description provided, for instance, by Sousa *et al*. (2021), and similar to other *Quercus* species, as, for instance *Q. petraea* (Gričar *et al*. 2015).

**Figure 3:**
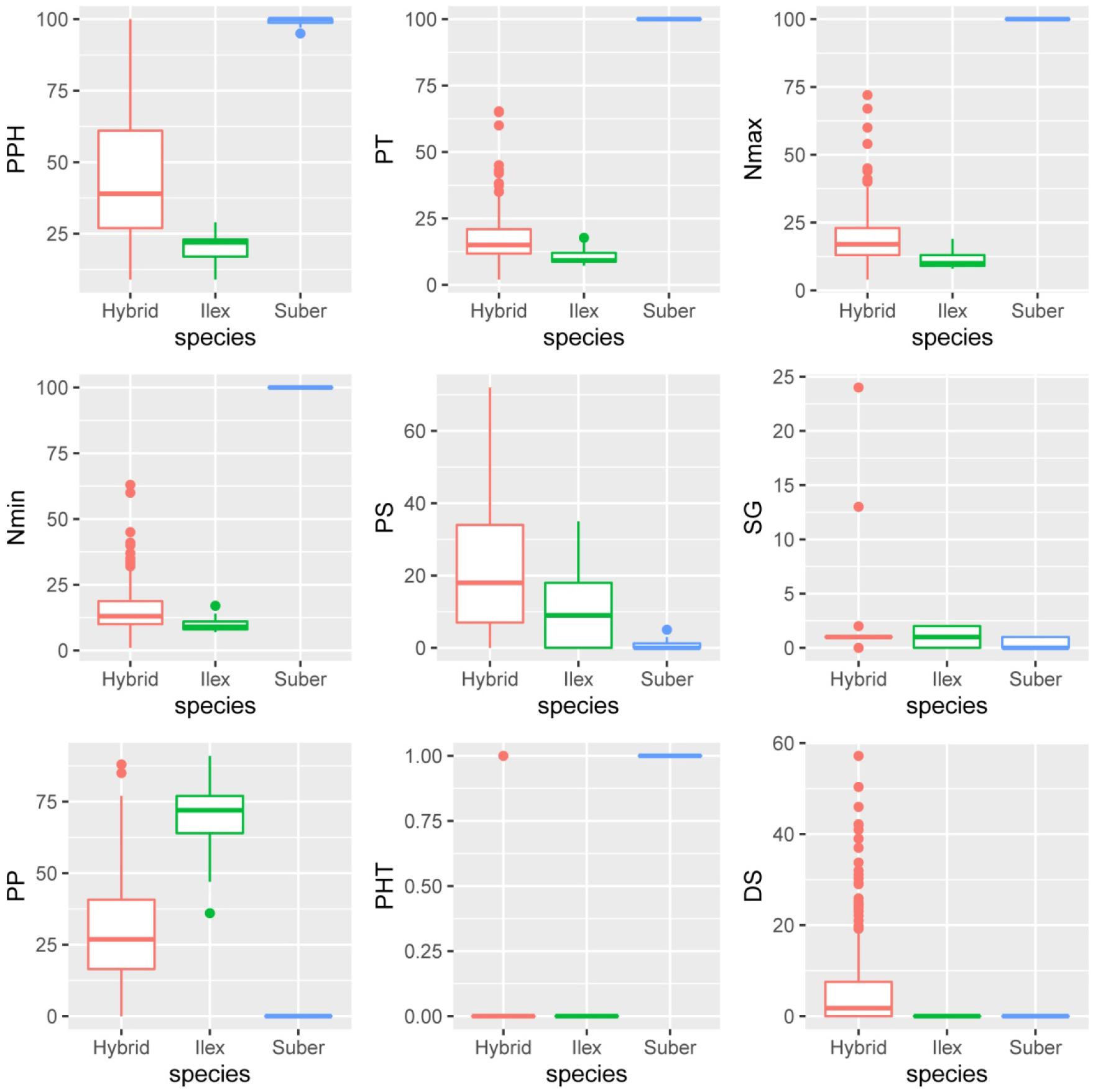
Boxplots of *Q. ilex, Q. suber* and the hybrids for all the variables related to outer bark anatomy described in **Table 1**.

ANOVA analyses show significant differences for all the quantitative variables regarding species and individuals (**Table 2)**, and so did Kruskal-Wallis test for the qualitative ones. Relationships among the variables were assessed by means of a correlation analysis (**Table S2**) Thus, phellem average thickness is highly correlated with maximum and minimum number of phellem cells and with the qualitative variable “phellogen type” (PHT), as expected, and negatively with the proportion of other tissues. This is consistent with Principal Component Analysis. The first two PC explain 80% of the total variance; the first one (59.5% of total variance) is mainly composed by variables related to phellem characteristics: phellem average thickness, maximum and minimum number of phellem cells and percentage of phellem tissue. Variables related to the other cell types contribute mainly to the second component (20.4%): percentages of parenchyma and, oppositely, percentages of schlerenchyma and suberized inactive phloem (although correlation of this suberization with the percentage of other tissues is not statistically significant) (**Figure 4**).

**Table 2:** Summary of the ANOVA results for the model expressed in eq. 1 and the quantitative variables described in Table 1. df: degrees of freedom; SS: sum of squares; MS: mean square; F: value of the F-test.

| **Variable** | **Factor** | **df** | **SS** | **MS** | **F** | **Pr (>F)** |
| --- | --- | --- | --- | --- | --- | --- |
| **PPH** | $\alpha_i$ (species) | 2 | 46228 | 23114 | 8.4 | 0.002 |
| | $\beta_{j(i)}$ (species:ind) | 20 | 89269 | 4463 | 16.9 | 0.001 |
| | $\gamma_{ijk}$ (residuals) | 368 | 97207 | 264 | | |
| **PT** | $\alpha_i$ (species) | 2 | 81547 | 40774 | 144.0 | 0.001 |
| | $\beta_{j(i)}$ (species:ind) | 20 | 8798 | 440 | 7.9 | 0.001 |
| | $\gamma_{ijk}$ (residuals) | 368 | 20525 | 56 | | |
| **Nmax** | $\alpha_i$ (species) | 2 | 78133 | 39066 | 129.6 | 0.001 |
| | $\beta_{j(i)}$ (species:ind) | 20 | 9342 | 467 | 7.6 | 0.001 |
| | $\gamma_{ijk}$ (residuals) | 368 | 22523 | 61 | | |
| **Nmin** | $\alpha_i$ (species) | 2 | 84678 | 42339 | 155.6 | 0.001 |
| | $\beta_{j(i)}$ (species:ind) | 20 | 8458 | 423 | 8.0 | 0.001 |
| | $\gamma_{ijk}$ (residuals) | 368 | 19549 | 53 | | |
| **PP** | $\alpha_i$ (species) | 2 | 36622 | 18311 | 17.3 | 0.001 |
| | $\beta_{j(i)}$ (species:ind) | 20 | 32934 | 1647 | 8.1 | 0.001 |
| | $\gamma_{ijk}$ (residuals) | 368 | 74518 | 202 | | |
| **PS** | $\alpha_i$ (species) | 2 | 5365 | 2682 | 2.3 | 0.119 |
| | $\beta_{j(i)}$ (species:ind) | 20 | 35762 | 1788 | 8.3 | 0.001 |
| | $\gamma_{ijk}$ (residuals) | 368 | 78966 | 215 | | |
| **DS** | $\alpha_i$ (species) | 2 | 948 | 473.8 | 1.5 | 0.242 |
| | $\beta_{j(i)}$ (species:ind) | 20 | 9757 | 487.8 | 7.9 | 0.001 |
| | $\gamma_{ijk}$ (residuals) | 368 | 22753 | 61.8 | | |

**Figure 4:**
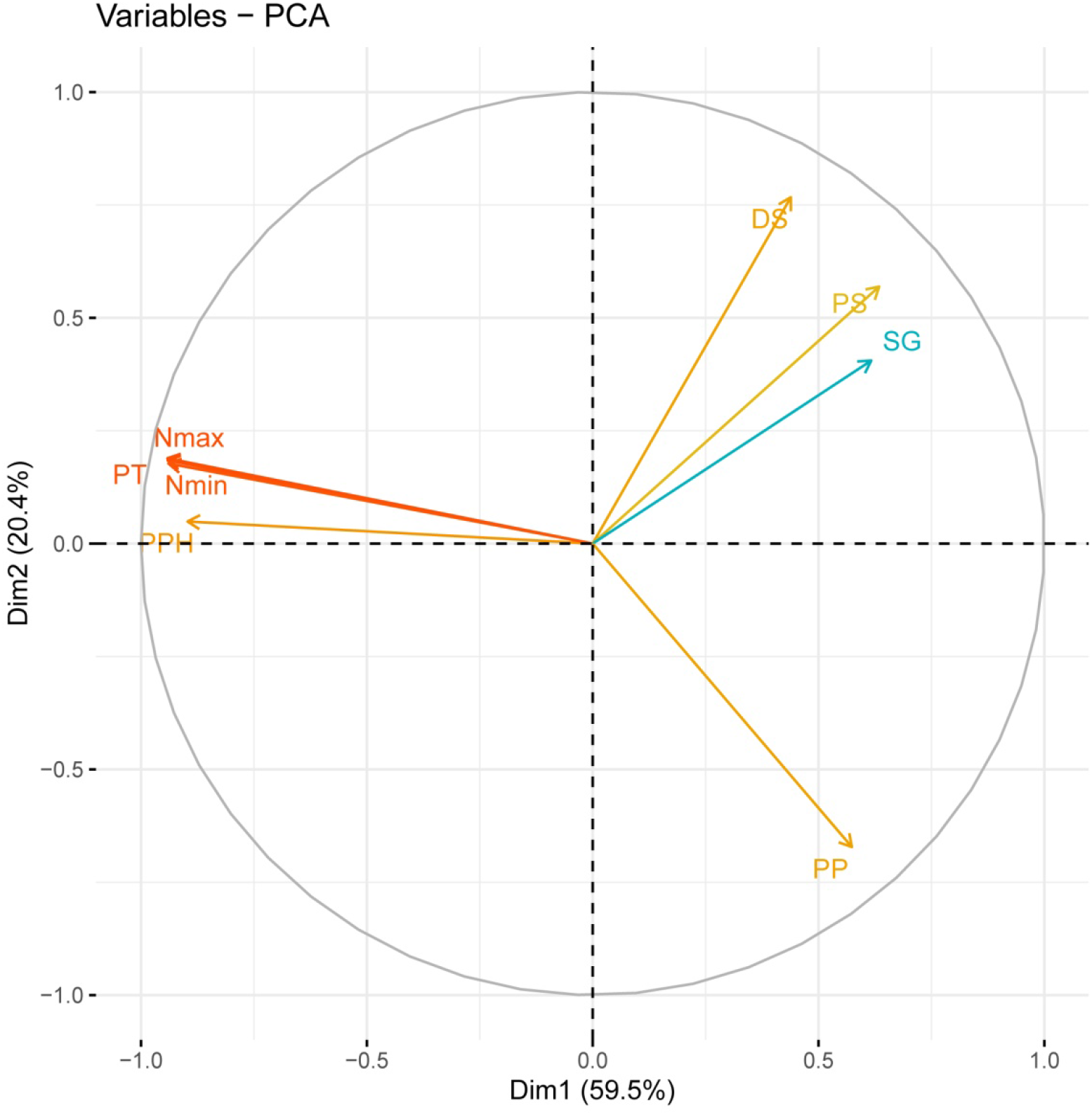
Relative contribution of the original variables to the first two PC for the multivariate analysis. The percentage of variance explained by each principal component is indicated.

The first component clearly discriminates *Q. suber* from *Q. ilex* and most hybrid individuals (**Figure 5**). However, certain hybrids, as FS01 and, to a minor extent, FS02 and FS18, are also separated from the rest of hybrids with this component. Notwithstanding average phellem thickness and maximum and minimum number of phellem cells in *Q. suber* is much higher than in these hybrids (and, therefore, cork oak samples appear far away in that axis), FS01, FS02 and FS18 have a much higher proportion of phellem in their barks than the other hybrids. This is specially the case of FS01, which presents almost 92% of the bark occupied by suberized ordered cells (“percentage of phellem”, PPH). This tissue also represents 67 and 64% of the bark in FS02 and FS18 samples, respectively, much larger than in the other hybrids. Average phellem thickness in these three hybrids is not that high, but in FS01 and FS02 (> 28 cells) it is 1.5-2.5 times thicker than in most other individuals. On the contrary, FS01 shows very low proportion of other cell types in its bark, and almost no suberization of remaining inactive phloem, as FS18.

**Figure 5:**
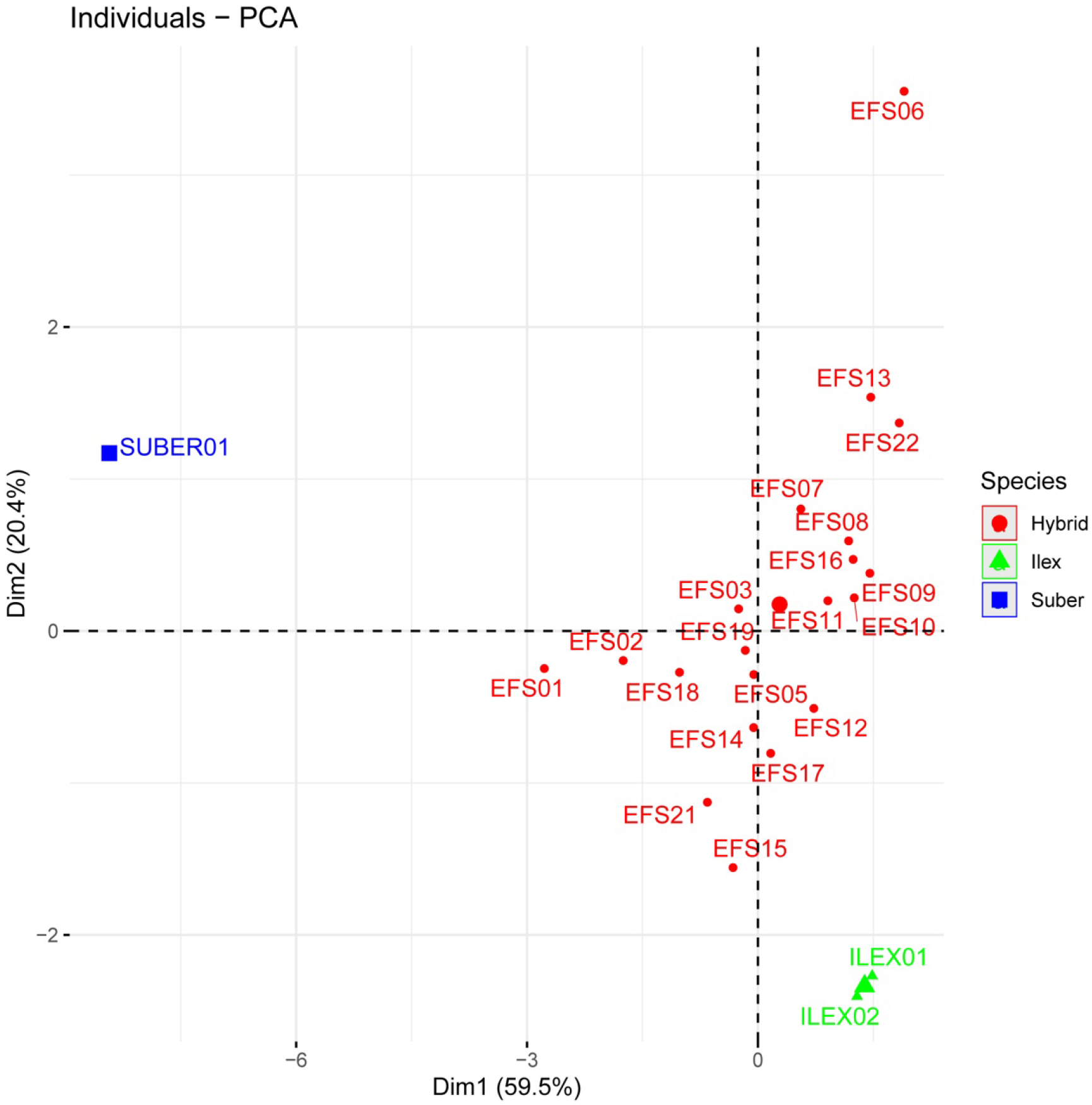
Biplot of the mean individual scores for the first two PCs for the multivariate analysis. The percentage of variance explained by each principal component is indicated.

The second component allows the discrimination of *Q. ilex* and hybrids and, among the latter, to distinguish hybrids with this unique phloem suberization from those more similar to *Q. ilex*. Thus, FS06 and FS22 show a large proportion of these cells (25% and 14% of their barks, respectively) (**Figure 2**). Interestingly, these individuals show the lowest proportion of phellem in their barks, and the thinnest periderms, close to *Q. ilex* values. The other variable contributing in the same sense to this component is the proportion of schlerenchymatic tissues in the bark, which reaches up to 42% in FS13 (also with >30% of phellem), and values above 30% in FS06, FS08, FS16 and FS22 (**Figure 6**).

**Figure 6:**
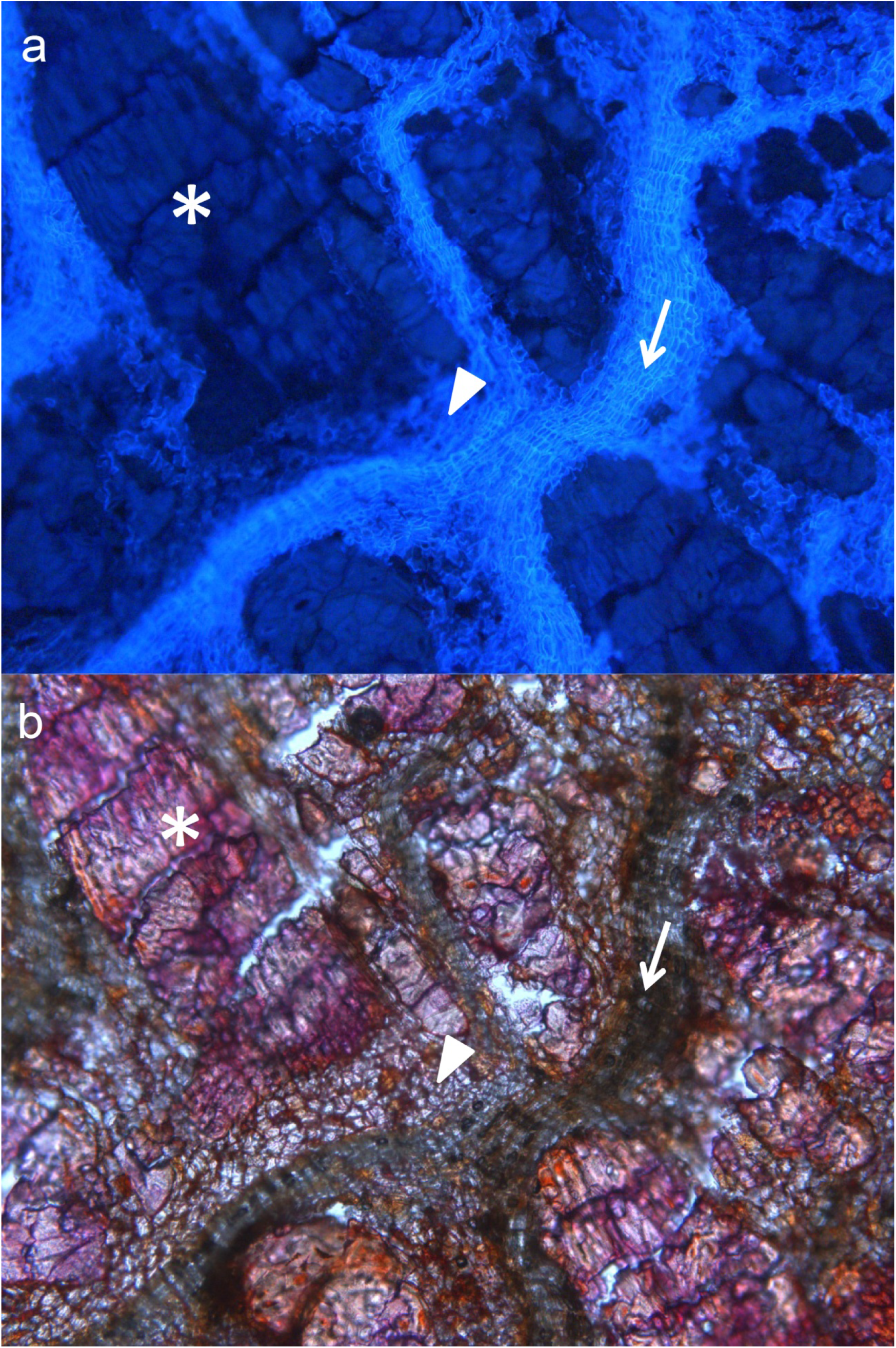
Cross section of the bark of a hybrid tree under UV (a) and visible light (b). Periderms (arrow), a large proportion of schlerenchymatic nodules (asterisk), and a lower proportion of suberized inactive phloem (arrowhead) can be appreciated.

On the contrary, the other individuals show slightly thicker periderms (30-50% of bark), and the largest proportions (more than 40% in some trees, and always >30%) of non-suberized parenchymatic tissue (inactive phellem), in their rhytidomes. This is the case, for instance, of FS15, FS17 and FS21, where inactive phloem suberization is very low (2.83, 1.15 and 0.12% of the bark) (**Figure 7**).

**Figure 7:**
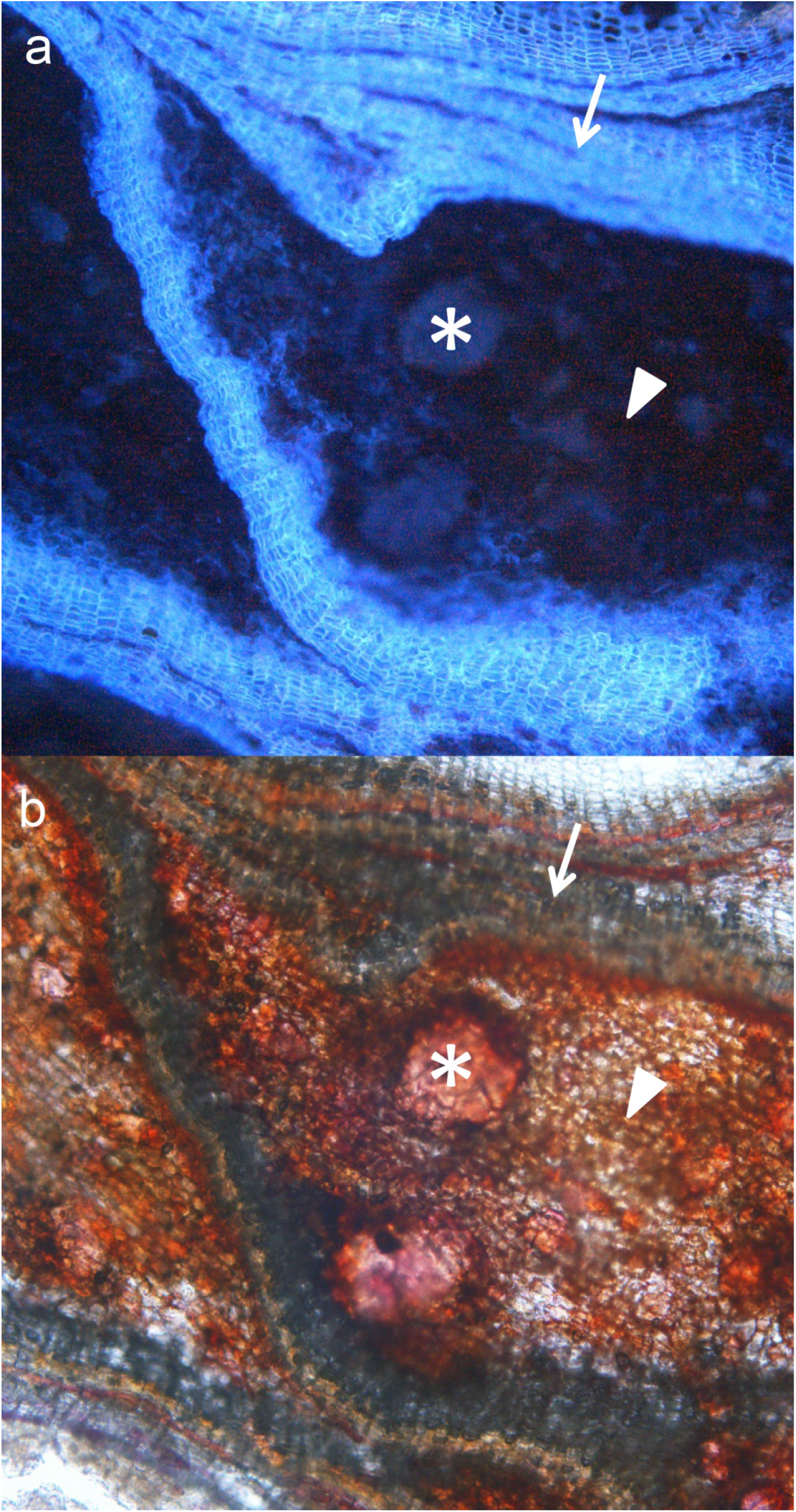
Cross section of the bark of a hybrid tree under UV (a) and visible light (b). Periderms (arrow), schlerenchymatic nodules (asterisk), and non-suberized inactive phloem (arrowhead) can be appreciated.

Other remarkable structures have been observed in the hybrids. For instance, schlerenchymatic nodules are often surrounded by thin periderms, as can be observed both in transverse and radial sections (**Figure 8**). These formations appear preferently in the outermost part of the bark, and could differentiate as an induced defense against a mechanical injure. The damaged area would be isolated by a traumatic periderm and then lignified, as described for conifers by Mullick (1975), Rittinger et al. (1987) or Wahlström (1992); alternatively, they could be formed constitutively. These observations are consistent with the “independent periderms” mentioned by Natividade (1936).

**Figure 8:**
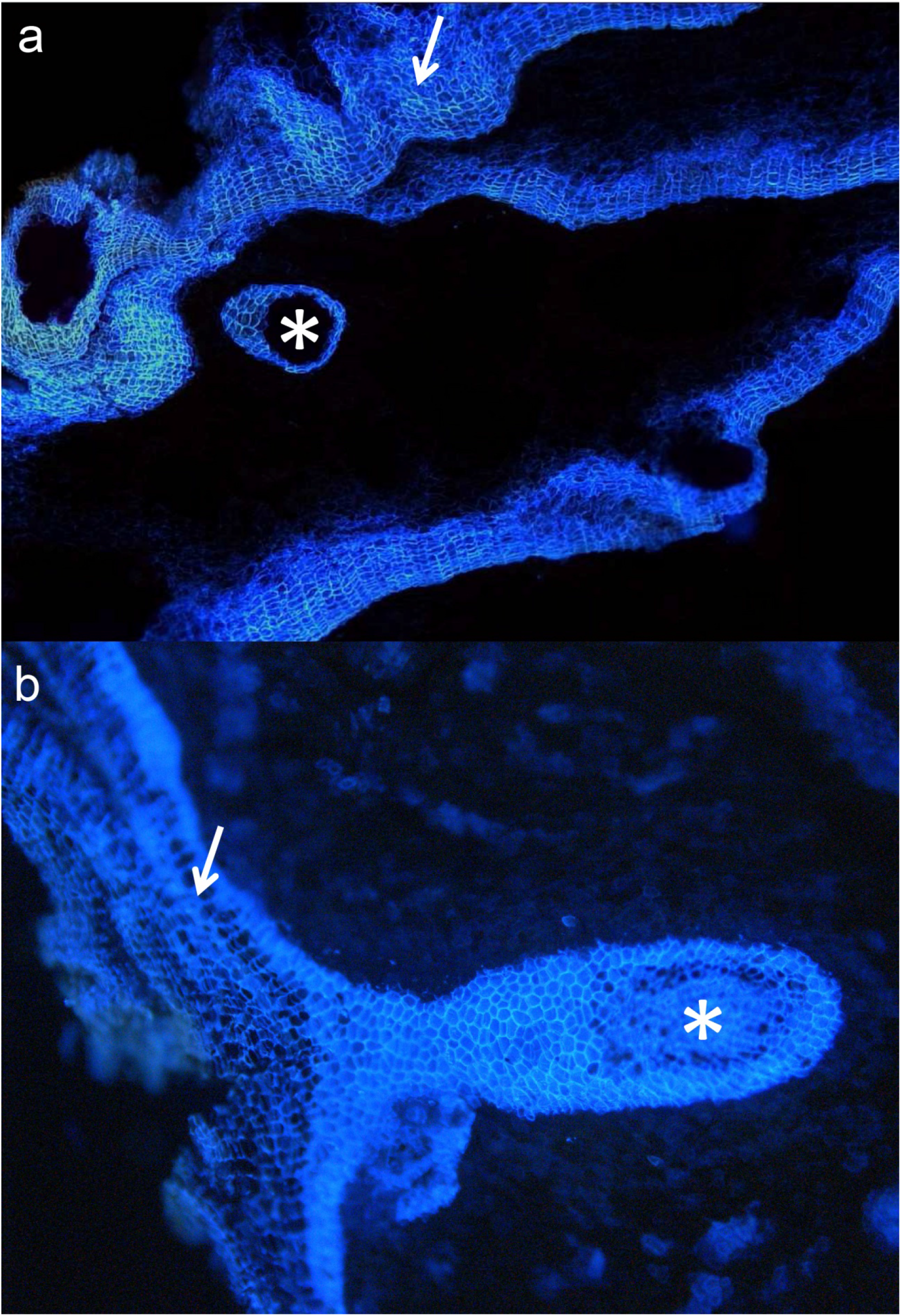
Cross section of the bark of a hybrid tree under UV light. **a**: A closed thin periderm surrounding a schlerenchymatic nodule (asterisk) can be appreciated. **b:** A similar structure, connected to the normal periderm (arrow).

Relationship between these bark anatomical variables and the contribution of each of the parental species to the genome of hybrid trees was also investigated. This contribution was estimated by López de Heredia *et al*. (2020), based on 9251 nuclear markers, mostly SNP. Nevertheless, only slight correlations were obtained (with R^2^ values below 20%), and were mostly due to the presence of FS01. This is the only individual classified as a probable backcross with *Q. suber*, with a 65-73.1% contribution of this species to its genome (q_s_). Consistently, this individual is characterized by a very large proportion of phellem in its bark (>90%) and an extremely low proportion of schlerenchyma (3%). Additionally, almost no suberization of the inactive phloem was detected (0.6% of the bark). If this individual is removed from the analysis, no significant relationship is detected for any of the variables. These other individuals are presumably first-generation hybrids, with estimated *Q. suber* genomic contributions (q_s_) between 45.4 and 52.6%. Lack of correlation is not surprising, first, due to the narrow range of q_s_ and, secondly, to its global character. This parameter provides an estimation of the average contribution of *Q. suber* to the whole genome, notwithstanding higher or lower contributions to specific genomic regions (not expected for F1 hybrids, though) and, more importantly, to the transcriptome, and consequently, to the proteome or metabolome.

### Concluding remarks

Development of a thick and corky bark, such as the one shown by *Q. suber*, relies on the presence of an active, long-living phellogen, able to produce enough anticlinal divisions in order to increase its circumference and keep the pace with the growth rhythm imposed by the activity of vascular cambium. Thus, the tangential tensions endured per each phellogen cell, which induce the differentiation of new successive inner phellogens in other speces, as *Q. ilex*, would remain moderate in *Q. suber*, and the periclinal divisions of this only phellogen, accumulated over the years, give rise to the characteristic thick phellem of this species.

All the hybrid trees analysed here but FS01 are presumably first generation hybrids, according to López de Heredia et al. (2020). Therefore, they are expected to carry a copy of all the genes underpinning the formation of a thick corky bark, as in *Q. suber*. Combination with *Q. ilex* genes hampers, however, the development of such a bark in those trees. Maybe trees need to have two copies of certain genes or alleles in order to produce corky bark. Indeed, the bark of FS01, the only individual classified as a backcross with *Q. suber* (and expected, therefore, to carry two *Q. suber* alleles in 50% of its loci, on average) is far more similar to cork oak bark. In addition, hybridization itself can induce epigenetic modifications, as pointed out by López de Heredia et al. (2020), which could affect the expression of such genes.

Thus, among F1 hybrids we have also detected relevant differences in their barks, regarding the number and thickness of their periderms, or the proportion of parenchymatic and sclerenchymatic cells in the rhytidome. Probably, the most striking feature is the generalized suberization of inactive phloem shown by several individuals. This feature also points out to the modification of the regulation pathways of suberin incorporation to the cell wall, induced by hybridization. Hybrid individuals appear, therefore, as an extremely interesting material for further research on the regulation of bark development, mainly by comparison with the parental species, *Q. suber* and *Q. ilex*.

## Acknowledgments

This work was funded by the projects AGL2015-67495-C2-2-R (Spanish Ministry of Economy and Competitiveness) and PID2019-110330GB-C22 (Spanish Ministry of Science and Innovation).

