## Supplementary material for "Qualitative and quantitative analysis of the constitutive bark of *Q. ilex* x *Q. suber* hybrids"

**Supplemental material**

**Table S1:** Mean and standard deviation values (among brackets) for the anatomical variables of the barks described in Table 1.

| species | ind | PPH | PHT | PT | Nmax | Nmin | DS | PS | SG | PP |
|---|---|---|---|---|---|---|---|---|---|---|
| Hyb | EFS01 | 91,67 (7,78) | 0,27 (0,46) | 28,49 (11,38) | 29,20 (11,05) | 27,73 (11,73) | 0,60 (1,3) | 3,00 (2,88) | 0,73 (0,46) | 4,73 (7,06) |
| Hyb | EFS02 | 67,04 (17,83) | 0 (0) | 28,35 (15,48) | 31,42 (16,54) | 25,83 (14,61) | 2,74 (7,57) | 10,54 (11,94) | 0,75 (0,61) | 19,68 (11,83) |
| Hyb | EFS03 | 59,25 (15,34) | 0 (0) | 16,25 (2,70) | 18,25 (3,35) | 14,60 (2,93) | 4,33 (5,54) | 15,60 (11,27) | 1,15 (0,59) | 20,82 (8,35) |
| Hyb | EFS05 | 40,08 (15,56) | 0 (0) | 21,11 (6,82) | 23,92 (7,76) | 18,08 (6,95) | 7,15 (9,91) | 14,75 (15,90) | 0,83 (0,72) | 38,02 (19,02) |
| Hyb | EFS06 | 27,75 (13,57) | 0 (0) | 15,15 (5,00) | 16,69 (5,02) | 13,63 (4,76) | 24,95 (14,30) | 32,38 (12,07) | 1,63 (0,50) | 14,93 (14,48) |
| Hyb | EFS07 | 42,32 (18,37) | 0 (0) | 16,80 (6,45) | 18,23 (6,33) | 15,27 (6,43) | 8,68 (13,06) | 22,95 (17,76) | 1,23 (0,61) | 26,05 (15,04) |
| Hyb | EFS08 | 36,25 (14,81) | 0 (0) | 11,03 (4,51) | 12,58 (4,50) | 9,58 (4,93) | 5,51 (5,08) | 31,17 (14,86) | 1,17 (0,72) | 27,07 (14,31) |
| Hyb | EFS09 | 36,26 (15,22) | 0 (0) | 13,32 (4,84) | 15,35 (6,07) | 11,39 (3,66) | 5,81 (6,45) | 18,17 (13,17) | 2,09 (4,82) | 39,75 (12,63) |
| Hyb | EFS10 | 40,18 (17,9) | 0 (0) | 12,75 (4,93) | 14,73 (4,90) | 11,18 (5,04) | 7,07 (11,15) | 12,91 (10,58) | 2,00 (3,69) | 39,84 (11,54) |
| Hyb | EFS11 | 30,89 (14,88) | 0 (0) | 16,18 (6,95) | 18,72 (7,58) | 14,22 (7,11) | 5,16 (4,83) | 27,83 (15,49) | 1,11 (0,32) | 36,12 (12,91) |
| Hyb | EFS12 | 32,47 (14,46) | 0 (0) | 13,48 (4,66) | 15,80 (4,81) | 11,73 (4,82) | 7,16 (8,75) | 18,47 (18,79) | 0,73 (0,59) | 41,90 (14,07) |
| Hyb | EFS13 | 29,00 (15,36) | 0 (0) | 13,78 (8,50) | 15,68 (8,74) | 12,04 (8,28) | 7,10 (6,56) | 42,36 (17,4) | 1,25 (0,52) | 21,55 (11,05) |
| Hyb | EFS14 | 44,70 (27,34) | 0 (0) | 16,02 (6,42) | 17,95 (7,11) | 14,55 (6,21) | 5,98 (13,40) | 14,00 (18,68) | 0,60 (0,60) | 35,32 (20,74) |
| Hyb | EFS15 | 43,90 (18,45) | 0 (0) | 17,52 (8,35) | 19,20 (8,57) | 16,00 (8,21) | 2,83 (5,84) | 8,50 (13,75) | 0,5 (0,71) | 44,77 (20,23) |
| Hyb | EFS16 | 32,10 (13,79) | 0 (0) | 12,23 (3,88) | 13,95 (4,27) | 10,55 (3,52) | 4,20 (5,59) | 33,65 (19,35) | 1,15 (0,37) | 30,05 (13,49) |
| Hyb | EFS17 | 49,46 (18,16) | 0 (0) | 12,14 (4,02) | 13,62 (4,41) | 10,62 (3,69) | 1,15 (2,98) | 17,54 (11,60) | 0,85 (0,38) | 31,85 (13,88) |
| Hyb | EFS18 | 64,47 (19,99) | 0 (0) | 19,73 (5,74) | 21,74 (5,99) | 17,58 (5,25) | 0 (0) | 19,21 (19,56) | 0,68 (0,48) | 16,32 (12,97) |
| Hyb | EFS19 | 44,13 (16,96) | 0 (0) | 20,81 (10,94) | 22,96 (11,01) | 19,00 (11,04) | 6,01 (7,74) | 16,87 (13,27) | 0,87 (0,55) | 32,99 (14,38) |
| Hyb | EFS21 | 52,30 (18,67) | 0 (0) | 19,15 (7,68) | 21,26 (7,96) | 17,09 (7,52) | 0,12 (0,58) | 13,96 (14,01) | 0,61 (0,5) | 33,62 (21,92) |
| Hyb | EFS22 | 24,44 (9,95) | 0 (0) | 10,93 (6,30) | 12,83 (6,39) | 9,28 (6,29) | 13,67 (11,60) | 30,50 (13,48) | 1,28 (0,57) | 31,38 (12,03) |
| *Q. ilex* | Ilex01 | 16,86 (7,01) | 0 (0) | 10,27 (3,33) | 11,14 (3,53) | 10,00 (3,16) | 0 (0) | 13,86 (11,02) | 1 (0,82) | 69,29 (15,35) |
| *Q. ilex* | Ilex02 | 21,40 (5,46) | 0 (0) | 10,95 (3,11) | 11,90 (3,51) | 10,10 (2,77) | 0 (0) | 8,90 (10,69) | 1,1 (0,88) | 69,70 (13,95) |
| *Q. suber* | Suber | 99,08 (1,62) | 1,00 (0) | 100 (0) | 100 (0) | 100 (0) | 0 (0) | 0,92 (1,62) | 0,33 (0,49) | 0 (0) |

**Table S2:** Pearson correlation estimate between the anatomical variables of the barks described in Table 1.

| | **PPH** | **PHT** | **PT** | **Nmax** | **Nmin** | **DS** | **PS** | **SG** | **PP** |
|---|---|---|---|---|---|---|---|---|---|
| **PPH** | 1 | | | | | | | | |
| **PHT** | 0,452 | 1,000 | | | | | | | |
| **PT** | 0,626 | 0,812 | 1,000 | | | | | | |
| **Nmax** | 0,631 | 0,793 | 0,994 | 1,000 | | | | | |
| **Nmin** | 0,615 | 0,825 | 0,996 | 0,984 | 1,000 | | | | |
| **DS** | -0,396 | -0,123 | -0,193 | -0,198 | -0,189 | 1,000 | | | |
| **PS** | -0,540 | -0,219 | -0,345 | -0,344 | -0,339 | 0,074 | 1,000 | | |
| **SG** | -0,143 | -0,095 | -0,130 | -0,131 | -0,125 | 0,022 | 0,194 | 1,000 | |
| **PP** | -0,583 | -0,314 | -0,385 | -0,389 | -0,379 | -0,046 | -0,264 | -0,006 | 1,000 |
